## Supplemental Information for "Genome-wide functional screens enable the prediction of high activity CRISPR-Cas9 and -Cas12a guides in *Yarrowia lipolytica*"

Figure S1: Design and validation of Cas12a sgRNA library for *Y. lipolytica.*
Figure S2: Replicate correlations at day 4 of the growth screen for Cas12a experiments.
Table S1: All replicate correlations for the Cas9 and Cas12a genome-wide growth screens.
Table S2: The twelve layers of DeepGuides' convolutional auto-encoder.
Table S3: The eleven layers in the second network of DeepGuide.
Table S4: Ablation analysis on Cas12a dataset.
Table S5: Ablation analysis on Cas9 dataset.
Figure S3: Genes used to validate DeepGuide and the observed phenotype of null mutants
Figure S4: Clustering of high and poor activity guides used for validation.
Figure S5: AUROC curves for all tools on Cas9 and Cas12a datasets.
Figure S6: Training and validation loss for DeepGuide with and without pre-training.
Table S6: Strains of yeast used in this study.
Table S7: Plasmids used or generated in this study.
Table S8: Primers used in this study.
Table S9: Transformation efficiencies achieved in the control and sample screens.
Table S10: Primers used for NGS fragment amplification.
Figure S7: Schematic and sequence information of Cas9 and Cas12a amplicons for NGS.
Table S11: Parameters for bioinformatics tools used in analysis of NGS reads.
Table S12: Correlation of SRA files names to demultiplexing information

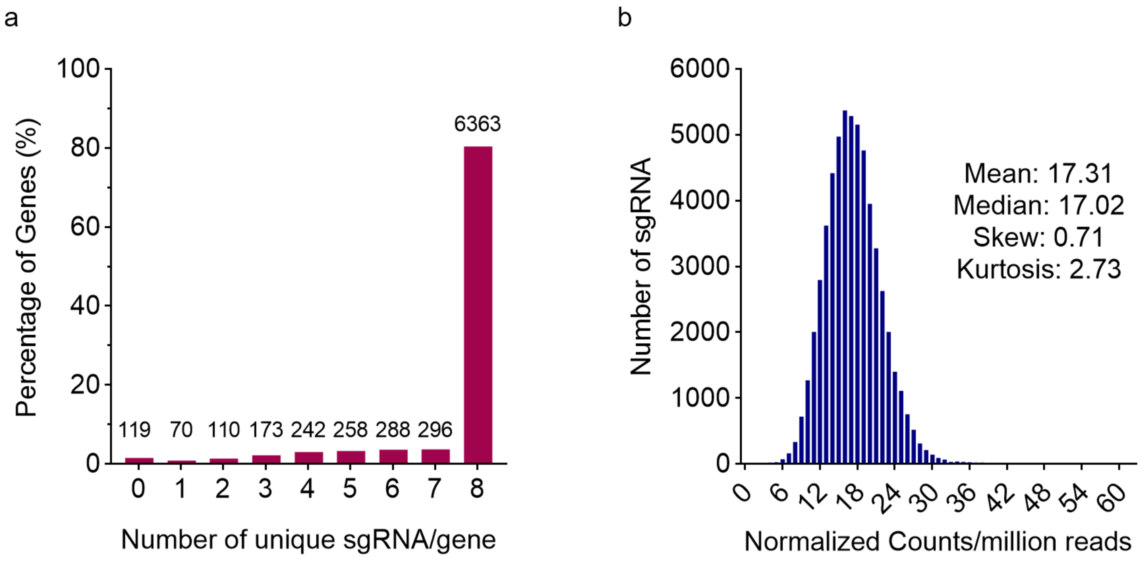


**Figure S1.** Design and validation of Cas12a and Cas9 sgRNA library for *Y. lipolytica* PO1f. (a) An 8-fold redundant sgRNA library was designed to target 7,919 protein coding genes in the *Y. lipolytica* CLIB89 strain, the parent strain of PO1f. Coding sequences were confirmed to be present in the PO1f genome sequence. Over 80% of the genes had 8 sgRNAs and over 91% of the genes had at least 5 sgRNAs. (b) A library consisting of 58,421 sgRNAs was synthesized by Agilent, cloned in-house and characterized by next generation sequencing. The library exhibited a tight normal distribution with nearly equal mean and median signifying minimal skew. The average representation of sgRNAs was ~100-fold (at 5.84 million reads which is 100 times the library size, we can calculate the mean representation of sgRNAs to be 5.84*17.31 = 101.09). **Note**: The Cas9 library design was previously reported in ref. 1 (see Figure S1). Additional details of this library are also provided in the materials and methods section of this manuscript.

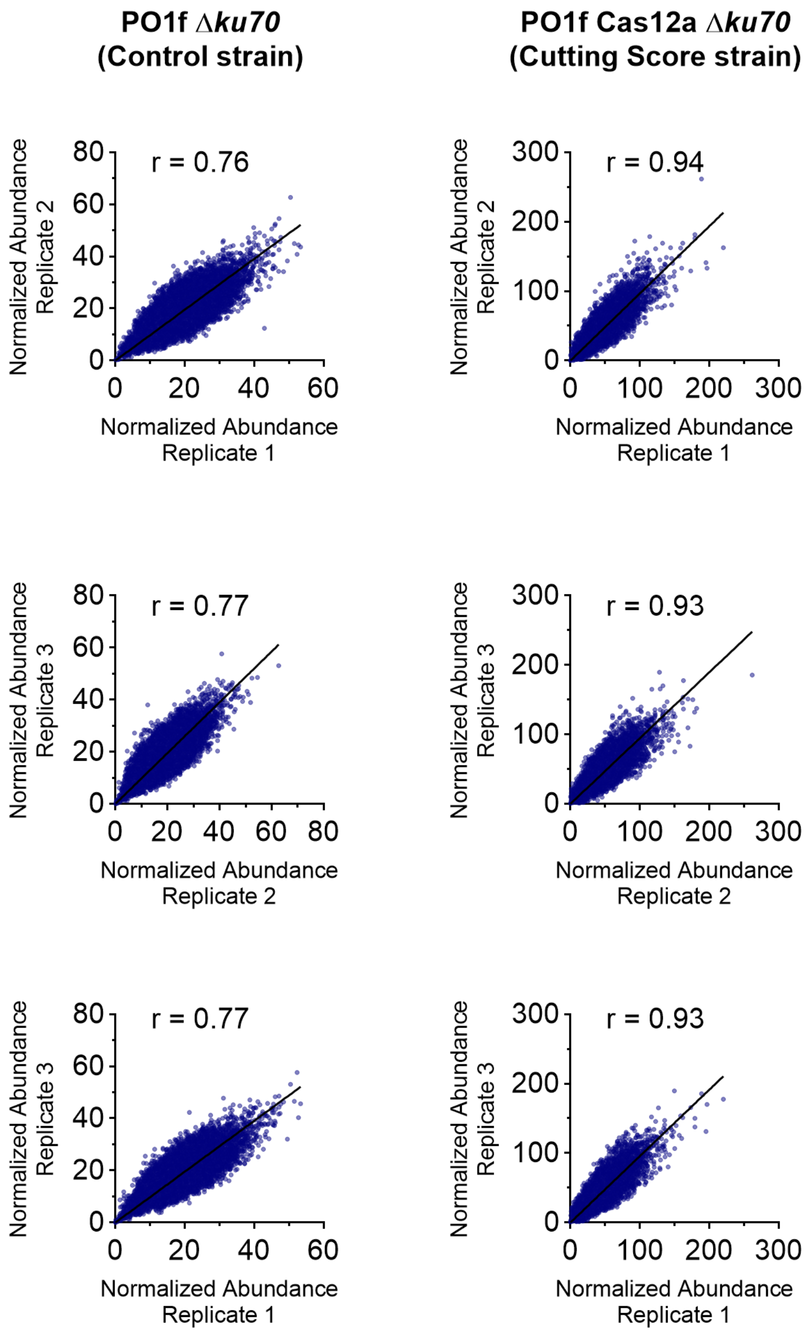


**Figure S2.** Replicate correlation graphs at Day 4 of the growth screen for Cas12a experiments. The column on the left shows pairwise correlations for the control strain while the column on the right shows the same for sample strain.

| Strain | Time point | Comparison | Pearson |
|---|---|---|---|
| PO1f *ku70* | Day 2 | 1 v. 2 | 0.765 |
| | | 1 v. 3 | 0.775 |
| | | 2 v. 3 | 0.738 |
| | Day 4 | 1 v. 2 | 0.756 |
| | | 1 v. 3 | 0.772 |
| | | 2 v. 3 | 0.762 |
| | Day 6 | 1 v. 2 | 0.797 |
| | | 1 v. 3 | 0.768 |
| | | 2 v. 3 | 0.799 |
| PO1f Cas12a *ku70* | Day 2 | 1 v. 2 | 0.902 |
| | | 1 v. 3 | 0.925 |
| | | 2 v. 3 | 0.892 |
| | Day 4 | 1 v. 2 | 0.936 |
| | | 1 v. 3 | 0.933 |
| | | 2 v. 3 | 0.927 |
| | Day 6 | 1 v. 2 | 0.918 |
| | | 1 v. 3 | 0.915 |
| | | 2 v. 3 | 0.905 |

| Strain | Time point | Comparison | Pearson |
|---|---|---|---|
| PO1f | Day 2 | 1 v. 2 | 0.988 |
| | | 1 v. 3 | 0.982 |
| | | 2 v. 3 | 0.980 |
| | Day 4 | 1 v. 2 | 0.829 |
| | | 1 v. 3 | 0.827 |
| | | 2 v. 3 | 0.858 |
| | Day 6 | 1 v. 2 | 0.818 |
| | | 1 v. 3 | 0.829 |
| | | 2 v. 3 | 0.855 |
| PO1f Cas9 *ku70* | Day 2 | 1 v. 2 | 0.972 |
| | | 1 v. 3 | 0.976 |
| | | 2 v. 3 | 0.972 |
| | Day 4 | 1 v. 2 | 0.886 |
| | | 1 v. 3 | 0.891 |
| | | 2 v. 3 | 0.973 |
| | Day 6 | 1 v. 2 | 0.877 |
| | | 1 v. 3 | 0.875 |
| | | 2 v. 3 | 0.968 |

**Table S1.** Replicate correlations for the genome-wide growth screens in *Y. lipolytica* with the Cas9 and Cas12a endonucleases. Cas9 data was previously reported in ref. [1] Note: Work conducted in ref. 1 uses PO1f with functional KU70 as the control strain.

| CAE (1st network) | Layer # | Layer type |
|---|---|---|
| Encoder | 1 | Convolution |
| | 2 | Batch Normalization |
| | 3 | Max Pooling |
| | 4 | Convolution |
| | 5 | Batch Normalization |
| | 6 | Average Pooling |
| Decoder | 7 | Up Sampling |
| | 8 | Batch Normalization |
| | 9 | Convolution |
| | 10 | Up Sampling |
| | 11 | Batch Normalization |
| | 12 | Convolution |

**Table S2.** The twelve layers in the convolutional auto-encoder (first network in DeepGuide); the autoencoder is composed by an encoder (layers 1-6) and a decoder (layers 7-12).

| 2nd network | Layer # | Layer type |
| --- | --- | --- |
| **Encoder** | 1 | Convolution |
| | 2 | Batch Normalization |
| | 3 | Max Pooling |
| | 4 | Convolution |
| | 5 | Batch Normalization |
| | 6 | Average Pooling |
| **Fully connected network** | 7 | Flatten |
| | 8 | Fully connected |
| | 9 | Fully connected |
| | 10 | Fully connected |
| | 11 | Multiplication |

**Table S3.** The eleven layers in the second network in DeepGuide, composed of an encoder (layers 1-6) and a fully connected network (layers 7-11).

| Cas12a | Layers | Spearman | Pearson |
|---|---|---|---|
| No pre-training (random weights), **no** back-propagation | encoder⇨flatten$_{7}$ | 0.060 | 0.070 |
| No pre-training (random weights), followed by back-propagation **only** on the flatten layer | encoder⇨flatten$_{7}$ | 0.451 | 0.455 |
| Pre-training of the encoder followed by back-propagation **only** the layers downstream of the encoder (flatten$_{7}$⇨…) | encoder⇨flatten$_{7}$ | 0.521 | 0.532 |
| | encoder⇨flatten$_{7}$⇨fc$_{8}$ | **0.527** | **0.534** |
| | encoder⇨flatten$_{7}$⇨fc$_{8}$⇨fc$_{9}$ | 0.505 | 0.517 |
| | encoder⇨flatten$_{7}$⇨fc$_{8}$⇨fc$_{9}$⇨fc$_{10}$ | 0.501 | 0.514 |
| | encoder⇨flatten$_{7}$⇨fc$_{8}$⇨fc$_{9}$⇨fc$_{10}$⇨mult$_{11}$ | 0.501 | 0.514 |
| Pre-training of the encoder followed by back-propagation on the entire network | encoder⇨flatten$_{7}$ | 0.637 | 0.641 |
| | encoder⇨flatten$_{7}$⇨fc$_{8}$ | 0.649 | 0.658 |
| | encoder⇨flatten$_{7}$⇨fc$_{8}$⇨fc$_{9}$ | **0.653** | **0.660** |
| | encoder⇨flatten$_{7}$⇨fc$_{8}$⇨fc$_{9}$⇨fc$_{10}$ | 0.653 | 0.660 |
| | encoder⇨flatten$_{7}$⇨fc$_{8}$⇨fc$_{9}$⇨fc$_{10}$⇨mult$_{11}$ | 0.653 | 0.660 |

**Table S4.** Ablation analysis on Cas12a dataset; green row (row 1) show the performance of the encoder (followed by a flatten layer) using random weights (no pre-training or backpropagation); purple row (row 2) show the performance of the encoder (followed by a flatten layer) using random weights and then performing back-propagation only on the flatten layer; blue rows (3-7) show the performance after pre-training the encoder and then running back-propagation only layers downstream of the encoder; pink rows (8-12) show the performance after pre-training and then running back-propagation on the whole network (including the encoder); correlation coefficients in bold corresponds to the best performance; fc = fully connected layer; pool = pooling layer; flatten = flatten layer; mult = multiplication layer (see Table S5 for the list of layers)

| Cas9 | Layers | Spearman r | Pearson r |
|---|---|---|---|
| No pre-training (random weights), **no** back-propagation | encoder⇨flatten$_7$ | 0.004 | 0.003 |
| No pre-training (random weights), followed by back-propagation **only** on the flatten layer | encoder⇨flatten$_7$ | 0.291 | 0.312 |
| Pre-training of the encoder followed by back-propagation **only** the layers downstream of the encoder (flatten$_7$⇨…) | encoder⇨flatten$_7$ | 0.316 | 0.353 |
| | encoder⇨flatten$_7$⇨fc$_8$ | 0.273 | 0.310 |
| | encoder⇨flatten$_7$⇨fc$_8$⇨fc$_9$ | 0.261 | 0.291 |
| | encoder⇨flatten$_7$⇨fc$_8$⇨fc$_9$⇨fc$_{10}$ | 0.269 | 0.305 |
| | encoder⇨flatten$_7$⇨fc$_8$⇨fc$_9$⇨fc$_{10}$⇨mult$_{11}$ | **0.345** | **0.388** |
| Pre-training of the encoder followed by back-propagation on the entire network | encoder⇨flatten$_7$ | 0.347 | 0.409 |
| | encoder⇨flatten$_7$⇨fc$_8$ | 0.364 | 0.424 |
| | encoder⇨flatten$_7$⇨fc$_8$⇨fc$_9$ | 0.357 | 0.414 |
| | encoder⇨flatten$_7$⇨fc$_8$⇨fc$_9$⇨fc$_{10}$ | 0.357 | 0.414 |
| | encoder⇨flatten$_7$⇨fc$_8$⇨fc$_9$⇨fc$_{10}$⇨mult$_{11}$ | **0.431** | **0.501** |

**Table S5.** Ablation analysis on Cas9 dataset; dataset; green row (row 1) show the performance of the encoder (followed by a flatten layer) using random weights (no pre-training or backpropagation); purple row (row 2) show the performance of the encoder (followed by a flatten layer) using random weights and then performing back-propagation only on the flatten layer; blue rows (3-7) show the performance after pre-training the encoder and then running back-propagation only layers downstream of the encoder; pink rows (8-12) show the performance after pre-training and then running back-propagation on the whole network (including the encoder); correlation coefficients in bold corresponds to the best performance; fc = fully connected layer; pool = pooling layer; flatten = flatten layer; mult = multiplication layer (see Table S5 for the list of layers)

| Gene Name | Function | Observed phenotype of null |
|---|---|---|
| MGA1 | Heat shock factor & pseudohyphal growth | Smooth colonies |
| RAS2 | GTP-binding protein, regulates filamentous growth | Smooth colonies |
| CAN1 | Arginine permease | Canavanine resistance |
| MFE1 | β-oxidation of long chain fatty acids | Oleic acid metabolism muted |

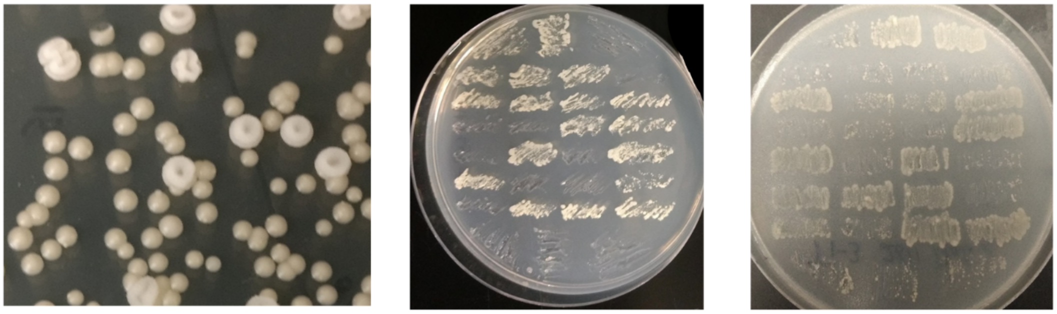

*ΔRAS2* & *ΔMGA1* *ΔCAN1* *ΔMFE1*

**Figure S3.** Genes selected for experimental validation of DeepGuide and the observed phenotype of the null mutants. MGA1 and RAS2 are implicated in the pseudohyphal and filamentous growth, and their null mutants show smooth colonies as shown in the picture on the left. CAN1 disruption confers resistance to L-Canavanine which is a toxic analog of Arginine. This leads to growth on plates supplemented with canavanine, as shown in the middle picture. MFE1 disruption renders *Y. lipolytica* unable to utilize oleic acid as a carbon source, and null mutants do not grow on plates with oleic acid as the sole carbon source as shown in the right-most picture.

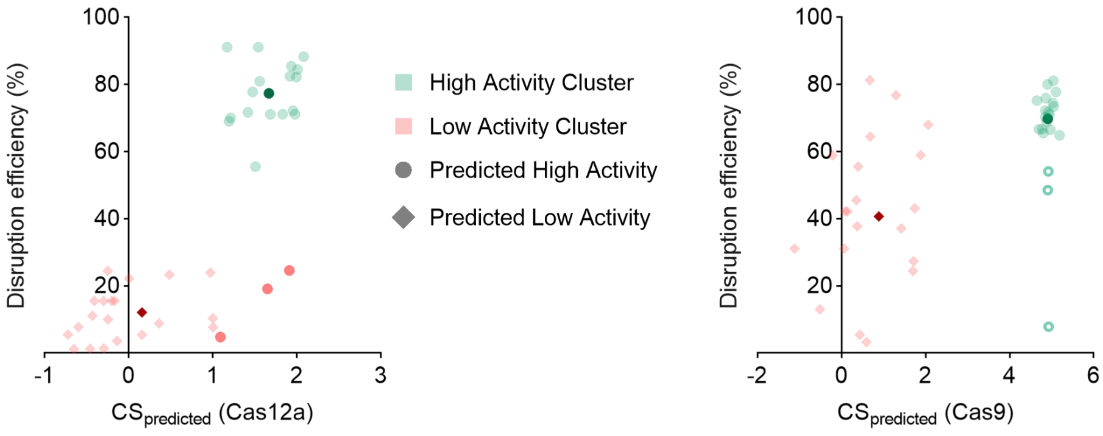


**Figure S4.** Clustering of high and poor activity guides used to validate DeepGuide. Predicted CS values and experimental disruption efficiencies for both the Cas12a and Cas9 were plotted on an XY scatter plot and a gaussian mixture model was used to cluster the sgRNA into two clusters (high and low activity). The high activity clusters are indicated in green, while the low activity clusters are indicated in red. Dark green and red points correspond to cluster centroids. Data point shape indicates whether the guide was predicted to be of high or low activity (circles are high activity, diamonds are low activity). Three predicted high activity guides cluster with low activity guides for Cas12a. For Cas9, three guides in the high activity cluster have a significantly higher euclidean distance from the cluster centroid and appear to be outliers (marked with empty circles).

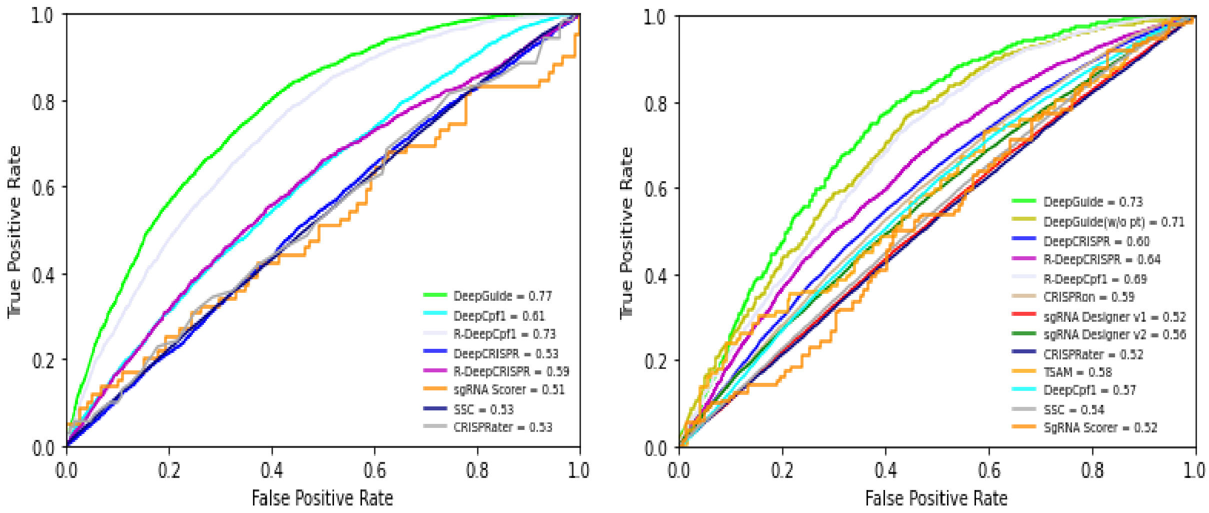


**Figure S5.** ROC plots and AUROC values for DeepGuide, DeepCpf1 (original and retrained), DeepCRISPR (original and retrained), sgRNA Scorer, SSC, and CRISPRater for the prediction of sgRNA activity on the Cas12a dataset (left) and the Cas9 dataset (right). DeepGuide had higher AUROC values than all other guide activity prediction algorithms. Guides with CS > 1.67 for Cas12a and CS > 4.91 for Cas9 were classified as active, and guides with a CS value below this threshold were classified as inactive.

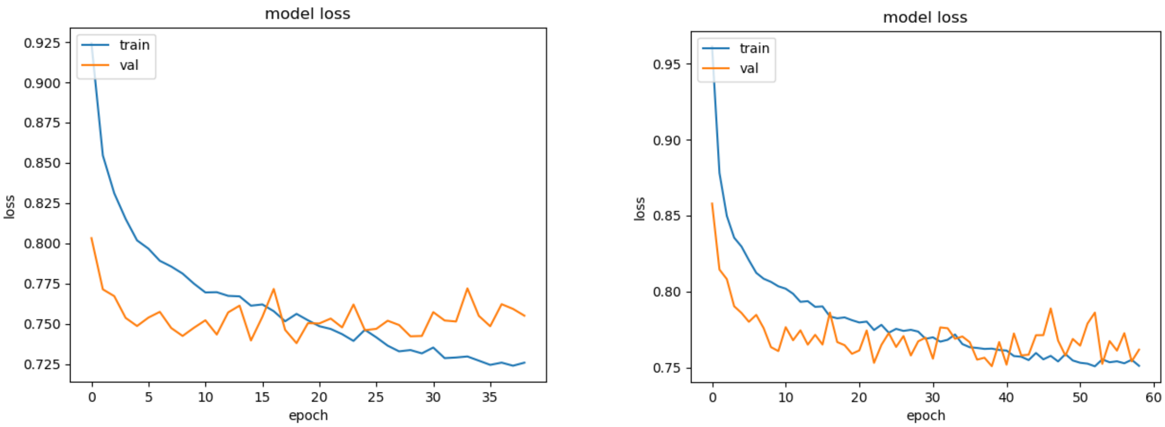


**Figure S6.** Training and validation loss for DeepGuide without pre-training (left) and with pre-training (right) as a function of the number of training epochs. These curves show that pre-training improves the architecture's generalization.

**Table S6.** Yeast strains used in this study.

| **Yeast strain genotype** | **Phenotype** |
|---|---|
| PO1f (MatA, *leu2-270*, *ura3-302*, *xpr2-322*, *axp-2*) | Wild type strain |
| PO1f *Δku70* | PO1f with disrupted KU70, which facilitates the non-homologous end joining DNA repair pathway |
| PO1f UAS1B8-TEF(136)-Cas9 -CycT::A08 | PO1f expressing *Y. lipolytica* codon optimized Cas9 gene at the A08 locus |
| PO1f UAS1B8-TEF(136)-LbCas12a -CycT::A08 | PO1f expressing *Y. lipolytica* codon optimized LbCas12a gene at the A08 locus |
| PO1f *Δku70* UAS1B8-TEF(136)-Cas9 -CycT::A08 | KU70 disrupted in Cas9 integrated PO1f strain |
| PO1f *Δku70* UAS1B8-TEF(136)-LbCas12a -CycT::A08 | KU70 disrupted in LbCas12a integrated PO1f strain |

**Table S7.** Plasmids used for genome wide CRISPR screens.

| Plasmid name | Reference | Function |
|---|---|---|
| pCpf1_yl | [2] | Plasmid for CRISPR-LbCas12a based gene editing in *Y. lipolytica* |
| pCRISPRyl | [3] | Plasmid for CRISPR-Cas9 based gene editing in *Y. lipolytica* |
| pLbCas12ayl | This study | Plasmid for CRISPR-LbCas12a based gene editing in *Y. lipolytic*a. sgRNA is flanked on either end by the direct repeat, to allow sgRNAs to end in T residues without being construed as part of the PolyT terminator |
| pHR_A08_hrGFP (Addgene #84615) | This study | Plasmid containing homology arms for integration of hrGFP into the A08 locus |
| pHR_A08_LbCas12a | This study | Plasmid containing homology arms for integration of LbCas12a into the A08 locus |
| pHR_A08_Cas9 | [1] | Plasmid containing homology arms for integration of Cas9 into the A08 locus |
| pLbCas12ayl-GW | This study | Vector containing sgRNA expression cassette for cloning Cas12a sgRNA library. (Does not contain Cas12a expression cassette) |
| pCas9yl-GW | [1] | Vector containing sgRNA expression cassette for cloning Cas9 sgRNA library. (Does not contain Cas9 expression cassette) |
| pCRISPRyl_KU70 | This study | CRISPR plasmid for the disruption of KU70 |

**Table S8.** Sequences of primers used in this study.

| Primer name | Primer Sequence |
|---|---|
| ExtraDR-F | CGGCGCAAATTTCTACTAAGTGTAGACTAGTAATTTCTACTAAGTGTAGATTTTTTACGTCTAAGAAACCATTATT |
| ExtraDR-R | AATAATGGTTTCTTAGACGTAAAAAATCTACACTTAGTAGAAATTACTAGTCTACACTTAGTAGAAATTTGCGCCG |
| Cpf1-Int-F | TGCCTGGAGCCGAGTACGGCATTGATTACTAGTCCGGGTTCGAAGGTACCAAG |
| Cpf1-Int-R | TTAGGCTGGGTCTCGAGAGCAAAGAAGCCTAGGGCAAATTAAAGCCTTCGAGCG |
| BRIDGE-F | CTAAATTTGATGAAAGGGGGATCCCCCGGGTGGCGTAATCATGGTCATAGCTGTTTCCTG |
| BRIDGE-R | CAGGAAACAGCTATGACCATGATTACGCCACCCGGGGGATCCCCCTTTCATCAAATTTAG |
| A08-Seq-F | AGCCGAGTACGGCATTGAT |
| A08-Seq-R | TCAATGTAGCCTCCTCCAACC |
| Tef_Seq-F | GTTGGGACTTTAGCCAAG |
| Lb1-R | CTTCTGCTTGGTCTTCTGGTTG |
| Lb2-F | AACCTGTACAACCAGAAGACCAAG |
| Lb3-F | AAGGAGACCAACCGAGACGAG |
| Lb4-F | AACCTGCACACCATGTACTTCAAG |
| Lb5-F | CCAGATCACCAACAAGTTCGAGTC |
| M13-F | GTAAAACGACGGCCAGT |
| InversePCR-F | TTTTTTACGTCTAAGAAACCATTATTATCATGACATTAACCT |
| InversePCR-R | TGCGCCGACCCGGAATCGAACCGGGGGCCC |
| OLS-F | GTTTAGTGGTAAAATCCATCGTTGCCATCG |
| OLS-R | GATACGCCTATTTTTATAGGTTAATGTCATG |
| qPCR-GW-F | TTATGAACTGAAAGTTGATGGC |
| qPCR-GW-R | TCACACAGGAAACAGCTATG |
| Cas9-RAS2-1 | TTCGATTCCGGGTCGGCGCACGCGGTCACTCCCCGCTCGTGTTTTAGAGCTAGAAATAGC |
| Cas9-RAS2-2 | TTCGATTCCGGGTCGGCGCACTCCACCAGTGGAGCCAACCGTTTTAGAGCTAGAAATAGC |
| Cas9-RAS2-3 | TTCGATTCCGGGTCGGCGCAACCTCCTGCAGCACCTCCAAGTTTTAGAGCTAGAAATAGC |
| Cas9-RAS2-4 | TTCGATTCCGGGTCGGCGCAGACTCTCAATGCTCCACCAGGTTTTAGAGCTAGAAATAGC |
| Cas9-RAS2-5 | TTCGATTCCGGGTCGGCGCAGATGTCGTAAACCAGAAGATGTTTTAGAGCTAGAAATAGC |
| Cas9-RAS2-6 | TTCGATTCCGGGTCGGCGCAAATCTAGGGCCTCCAAAGACGTTTTAGAGCTAGAAATAGC |
| Cas9-RAS2-7 | TTCGATTCCGGGTCGGCGCATCCCGTTCCTGTGGTTAGTAGTTTTAGAGCTAGAAATAGC |

| | |
|---|---|
| Cas9-RAS2-8 | TTCGATTCCGGGTCGGCGCATGTTGGAGTCGACCTGGAAGG TTTTAGAGCTAGAAATAGC |
| Cas9-RAS2-9 | TTCGATTCCGGGTCGGCGCAAAGCTGTGGGTGCACTGGTCG TTTTAGAGCTAGAAATAGC |
| Cas9-RAS2-10 | TTCGATTCCGGGTCGGCGCAGGAACCAGAGGACTAAGCTG GTTTTAGAGCTAGAAATAGC |
| Cas9-MGA1-1 | TTCGATTCCGGGTCGGCGCACTGTTGCGCGGCCTGGGTCGG TTTTAGAGCTAGAAATAGC |
| Cas9-MGA1-2 | TTCGATTCCGGGTCGGCGCAACTGGCCAAGGAGCCTGCTGG TTTTAGAGCTAGAAATAGC |
| Cas9-MGA1-3 | TTCGATTCCGGGTCGGCGCATTGCGGCAGAGGCATGGTTTG TTTTAGAGCTAGAAATAGC |
| Cas9-MGA1-4 | TTCGATTCCGGGTCGGCGCACAGAGGCATGGTTTCGGCGCG TTTTAGAGCTAGAAATAGC |
| Cas9-MGA1-5 | TTCGATTCCGGGTCGGCGCAGCCCGGCGAGGAGTTCTCCAG TTTTAGAGCTAGAAATAGC |
| Cas9-MGA1-6 | TTCGATTCCGGGTCGGCGCAAAGACGGAGTTTGTGGGTGGG TTTTAGAGCTAGAAATAGC |
| Cas9-MGA1-7 | TTCGATTCCGGGTCGGCGCAAGAGAGACAGTGTGCCCTTGG TTTTAGAGCTAGAAATAGC |
| Cas9-MGA1-8 | TTCGATTCCGGGTCGGCGCAGTAGGGGGCGCCTGTCCGTCG TTTTAGAGCTAGAAATAGC |
| Cas9-MGA1-9 | TTCGATTCCGGGTCGGCGCAGAGTGTGGTGGCGGAGTAGA GTTTTAGAGCTAGAAATAGC |
| Cas9-MGA1-10 | TTCGATTCCGGGTCGGCGCATGCGCGGCCTGGGTCGTGGGG TTTTAGAGCTAGAAATAGC |
| Cas9-CAN1-1 | TTCGATTCCGGGTCGGCGCATCAAACGATTACCCACCCTCGT TTTAGAGCTAGAAATAGC |
| Cas9-CAN1-2 | TTCGATTCCGGGTCGGCGCATTACCCACCCTCCGGGACTGG TTTTAGAGCTAGAAATAGC |
| Cas9-CAN1-3 | TTCGATTCCGGGTCGGCGCACCACATCCACATCAACCACAG TTTTAGAGCTAGAAATAGC |
| Cas9-CAN1-4 | TTCGATTCCGGGTCGGCGCACATCAACCACACGGCCCACTG TTTTAGAGCTAGAAATAGC |
| Cas9-CAN1-5 | TTCGATTCCGGGTCGGCGCACACCAGTGGCCACGACCTGGG TTTTAGAGCTAGAAATAGC |
| Cas9-CAN1-6 | TTCGATTCCGGGTCGGCGCAAGTGGGCCGTGTGGTTGATGG TTTTAGAGCTAGAAATAGC |
| Cas9-CAN1-7 | TTCGATTCCGGGTCGGCGCACCGTGTGGTTGATGTGGATGG TTTTAGAGCTAGAAATAGC |
| Cas9-CAN1-8 | TTCGATTCCGGGTCGGCGCAGTGGATGTGGGCCTCAGTCCG TTTTAGAGCTAGAAATAGC |
| Cas9-CAN1-9 | TTCGATTCCGGGTCGGCGCAGATGTGGGCCTCAGTCCCGGG TTTTAGAGCTAGAAATAGC |

| | |
|---|---|
| Cas9-CAN1-10 | TTCGATTCCGGGTCGGCGCATGGGCCTCAGTCCCGGAGGGG TTTTAGAGCTAGAAATAGC |
| Cas9-MFE1-1 | TTCGATTCCGGGTCGGCGCATGGTGAGACCCTGAAGGTTGG TTTTAGAGCTAGAAATAGC |
| Cas9-MFE1-2 | TTCGATTCCGGGTCGGCGCAGGTGTTATCCCTTACATGGGGT TTTAGAGCTAGAAATAGC |
| Cas9-MFE1-3 | TTCGATTCCGGGTCGGCGCACGTACTTCTGCTTAAGGAAGG TTTTAGAGCTAGAAATAGC |
| Cas9-MFE1-4 | TTCGATTCCGGGTCGGCGCAGACAAGATCCCAGTCCTTGTG TTTTAGAGCTAGAAATAGC |
| Cas9-MFE1-5 | TTCGATTCCGGGTCGGCGCAATACTTGAGCTCATTAGCCTGT TTTAGAGCTAGAAATAGC |
| Cas9-MFE1-6 | TTCGATTCCGGGTCGGCGCACTGCTTTCGGAAGTAAGGCCG TTTTAGAGCTAGAAATAGC |
| Cas9-MFE1-7 | TTCGATTCCGGGTCGGCGCAAAAGCAGGGTCGATGTGAAG GTTTTAGAGCTAGAAATAGC |
| Cas9-MFE1-8 | TTCGATTCCGGGTCGGCGCAGTCGATGAAATTAAGGCCCTG TTTTAGAGCTAGAAATAGC |
| Cas9-MFE1-9 | TTCGATTCCGGGTCGGCGCAGTTGTTGTCAACGATCTTGGG TTTTAGAGCTAGAAATAGC |
| Cas9-MFE1-10 | TTCGATTCCGGGTCGGCGCACTTGGATCGGACAGACTCGAG TTTTAGAGCTAGAAATAGC |
| Cas12a-RAS2-1 | TTTCTACTAAGTGTAGATGAGGCCCTAGATTACTTCAACGAC AAATTTCTACTAAGTGTA |
| Cas12a-RAS2-2 | TTTCTACTAAGTGTAGATGACCACCTAACGACGCGAAAAAA CAAATTTCTACTAAGTGTA |
| Cas12a-RAS2-3 | TTTCTACTAAGTGTAGATCGACATCACAGCCCCCCAGTCTTT GAATTTCTACTAAGTGTA |
| Cas12a-RAS2-4 | TTTCTACTAAGTGTAGATGGCACCCGCACACCGGCCCCAGC TTAATTTCTACTAAGTGTA |
| Cas12a-RAS2-5 | TTTCTACTAAGTGTAGATCATGAATCCGCATCCATGCTCGCG CAATTTCTACTAAGTGTA |
| Cas12a-RAS2-6 | TTTCTACTAAGTGTAGATCATTGTCATTCTTGGAGAGGGAGG TAATTTCTACTAAGTGTA |
| Cas12a-RAS2-7 | TTTCTACTAAGTGTAGATCGTCGCGACTGGGTGTGTCTGATC GAATTTCTACTAAGTGTA |
| Cas12a-RAS2-8 | TTTCTACTAAGTGTAGATGCGTCGTTAGGTGGTCCAAAACG AGAATTTCTACTAAGTGTA |
| Cas12a-RAS2-9 | TTTCTACTAAGTGTAGATCTGAAGTTTCCATGAATCCGCATC CAATTTCTACTAAGTGTA |
| Cas12a-RAS2-10 | TTTCTACTAAGTGTAGATCGCGACTTTGCGCACTATAGATGA GAATTTCTACTAAGTGTA |
| Cas12a-MGA1-1 | TTTCTACTAAGTGTAGATTGGGTGGTGGATTCGCTGAAGCGC TAATTTCTACTAAGTGTA |

| | |
|---|---|
| Cas12a-MGA1-2 | TTTCTACTAAGTGTAGATATGGTCTGCGTCCAACGACTCGTTCAATTTCTACTAAGTGTA |
| Cas12a-MGA1-3 | TTTCTACTAAGTGTAGATGGCGGCATGTGCTCGACCCGTTCTTAATTTCTACTAAGTGTA |
| Cas12a-MGA1-4 | TTTCTACTAAGTGTAGATTGCGCCAGCTCAACATGTACGGCTTAATTTCTACTAAGTGTA |
| Cas12a-MGA1-5 | TTTCTACTAAGTGTAGATGGTGGCCCATGGCGTGTGCCACCCGAATTTCTACTAAGTGTA |
| Cas12a-MGA1-6 | TTTCTACTAAGTGTAGATTCAACAATCTGCAGCAGCGTCTGCAAATTTCTACTAAGTGTA |
| Cas12a-MGA1-7 | TTTCTACTAAGTGTAGATTTGAACCCAGAAGGGGGCGACAAGAAATTTCTACTAAGTGTA |
| Cas12a-MGA1-8 | TTTCTACTAAGTGTAGATGAGTGGTGCCGGGCTTCTTGTTATCTTTTTTACGTCTAAGAA |
| Cas12a-MGA1-9 | TTTCTACTAAGTGTAGATCCTGCTGGATGTCCTCCCGCGAATCAATTTCTACTAAGTGTA |
| Cas12a-MGA1-10 | TTTCTACTAAGTGTAGATGGCGCCGGAGGCTGTGTGGCGACGGAATTTCTACTAAGTGTA |
| Cas12a-CAN1-1 | TTTCTACTAAGTGTAGATCTACCCGATATCTGTCACAGTCGTTAATTTCTACTAAGTGTA |
| Cas12a-CAN1-2 | TTTCTACTAAGTGTAGATACGACCCCAAGCTGACCGATGACTCAATTTCTACTAAGTGTA |
| Cas12a-CAN1-3 | TTTCTACTAAGTGTAGATGGCAGGAAACTCCAACGTCTACATTAATTTCTACTAAGTGTA |
| Cas12a-CAN1-4 | TTTCTACTAAGTGTAGATGTCTGCTGGCCTTCATGTCTGTGTCAATTTCTACTAAGTGTA |
| Cas12a-CAN1-5 | TTTCTACTAAGTGTAGATGTGCCTCCATGGGCTGGCTATACTGAATTTCTACTAAGTGTA |
| Cas12a-CAN1-6 | TTTCTACTAAGTGTAGATCATCTTCTACATTGGCTCTATCTTCAATTTCTACTAAGTGTA |
| Cas12a-CAN1-7 | TTTCTACTAAGTGTAGATTGGGGTTCTGGGCCTCACCGGCAGTAATTTCTACTAAGTGTA |
| Cas12a-CAN1-8 | TTTCTACTAAGTGTAGATCTTGTGCGAGGGCACCTCCTCTGAGTTTTTTACGTCTAAGAA |
| Cas12a-CAN1-9 | TTTCTACTAAGTGTAGATGTGCGGTTCCGGAGTCAGCCAGGGCAATTTCTACTAAGTGTA |
| Cas12a-CAN1-10 | TTTCTACTAAGTGTAGATCTCGAATTTGCATCTTCTACATTGGAATTTCTACTAAGTGTA |
| Cas12a-MFE1-1 | TTTCTACTAAGTGTAGATAGAGCCCCACCTACCCTAACGGCCCAATTTCTACTAAGTGTA |
| Cas12a-MFE1-2 | TTTCTACTAAGTGTAGATGCCATGTAACCAGCACCGACCTCGTAATTTCTACTAAGTGTA |
| Cas12a-MFE1-3 | TTTCTACTAAGTGTAGATGGGGGTGACACCCTTCTTGGTGTTGAATTTCTACTAAGTGTA |

| | |
|---|---|
| Cas12a-MFE1-4 | TTTCTACTAAGTGTAGATGGTGCCTACAAGGTTACCCGAGCTGAATTTCTACTAAGTGTA |
| Cas12a-MFE1-5 | TTTCTACTAAGTGTAGATATGTCCACCTCAACGGTACTTACTCAATTTCTACTAAGTGTA |
| Cas12a-MFE1-6 | TTTCTACTAAGTGTAGATCCGACTTTCTGGTGATTACAACCCTAATTTCTACTAAGTGTA |
| Cas12a-MFE1-7 | TTTCTACTAAGTGTAGATCGGAAACTTCGGCCAGACCAACTACAATTTCTACTAAGTGTA |
| Cas12a-MFE1-8 | TTTCTACTAAGTGTAGATGGTCGTTTCGCTTCGCTGCGCTTGTAATTTCTACTAAGTGTA |
| Cas12a-MFE1-9 | TTTCTACTAAGTGTAGATAAGAAGTCAGCAGGGCCGTTAGGGTAATTTCTACTAAGTGTA |
| Cas12a-MFE1-10 | TTTCTACTAAGTGTAGATTCCTTCTGTGTGGTGTCGTTTTGGGAATTTCTACTAAGTGTA |

**Table S9.** Transformation efficiencies measured as x$10^6$ transformants, for all replicates in the control and treatment strains.

| ***Strain*** | ***Replicate Transformation Efficiency (x$10^6$ transformants)*** | | |
|---|---|---|---|
| | **R1** | **R2** | **R3** |
| PO1f *Δku70* | 689 | 621 | 543 |
| PO1f Cas12a *Δku70* | 506 | 429 | 441 |

**Table S10.** Primers used for NGS fragment amplification

| Primer name | Primer Sequence | Illumina Barcode (Reverse primer) / Pseudo-Barcode (Forward primer) for demultiplexing |
|---|---|---|
| ILU1-F | AATGATACGGCGACCACCGAGATCTACACTCTTTCCCTACACGACGCTCTTCCGATCTTTCCGGGTCGGCGCAAATTTC | ^TTCCGG |
| ILU2-F | AATGATACGGCGACCACCGAGATCTACACTCTTTCCCTACACGACGCTCTTCCGATCTAGATCGGGTCGGCGCAAATTTCT | ^AGATCG |
| ILU3-F | AATGATACGGCGACCACCGAGATCTACACTCTTTCCCTACACGACGCTCTTCCGATCTGCTATTCGGGTCGGCGCAAATTTCT | ^GCTATT |
| ILU4-F | AATGATACGGCGACCACCGAGATCTACACTCTTTCCCTACACGACGCTCTTCCGATCTCAGGACTACGGGTCGGCGCAAATTTCT | ^CAGGAC |
| ILU1-R | CAAGCAGAAGACGGCATACGAGATTCGCCTTGGTGACTGGAGTTCAGACGTGTGCTCTTCCGATCTTAGAGGATCTGGGCCTCGTGATAC | CAAGGCGA |
| ILU2-R | CAAGCAGAAGACGGCATACGAGATGACGAGAGGTGACTGGAGTTCAGACGTGTGCTCTTCCGATCTTAGAGGATCTGGGCCTCGTGATAC | CTCTCGTC |
| ILU3-R | CAAGCAGAAGACGGCATACGAGATAGACTTGGGTGACTGGAGTTCAGACGTGTGCTCTTCCGATCTTAGAGGATCTGGGCCTCGTGATAC | CCAAGTCT |
| ILU4-R | CAAGCAGAAGACGGCATACGAGATCTGTATTAGTGACTGGAGTTCAGACGTGTGCTCTTCCGATCTTAGAGGATCTGGGCCTCGTGATAC | TAATACAG |
| ILU5-R | CAAGCAGAAGACGGCATACGAGATCCTGAACCGTGACTGGAGTTCAGACGTGTGCTCTTCCGATCTTAGAGGATCTGGGCCTCGTGATAC | GGTTCAGG |
| ILU6-R | CAAGCAGAAGACGGCATACGAGATATCAGGTTGTGACTGGAGTTCAGACGTGTGCTCTTCCGATCTTAGAGGATCTGGGCCTCGTGATAC | AACCTGAT |
| ILU7-R | CAAGCAGAAGACGGCATACGAGATTAGGTGACGTGACTGGAGTTCAGACGTGTGCT | GTCACCTA |

| | | |
|---|---|---|
| | CTTCCGATCTTAGAGGATCTGGGCCTCGT GATAC | |
| ILU8-R | CAAGCAGAAGACGGCATACGAGATCGAA CAGTGTGACTGGAGTTCAGACGTGTGCT CTTCCGATCTTAGAGGATCTGGGCCTCGT GATAC | ACTGTTCG |
| ILU9-R | CAAGCAGAAGACGGCATACGAGATGTTC GATCGTGACTGGAGTTCAGACGTGTGCTC TTCCGATCTTAGAGGATCTGGGCCTCGTG ATAC | GATCGAAC |
| ILU10-R | CAAGCAGAAGACGGCATACGAGATACCT AGCTGTGACTGGAGTTCAGACGTGTGCC TTCCGATCTTAGAGGATCTGGGCCTCGTG ATAC | AGCTAGGT |
| ILU11-R | CAAGCAGAAGACGGCATACGAGATAGAG ATGAGTGACTGGAGTTCAGACGTGTGCTC TTCCGATCTTAGAGGATCTGGGCCTCGTG ATAC | TCATCTCT |
| ILU12-R | CAAGCAGAAGACGGCATACGAGATCTGG ACTTGTGACTGGAGTTCAGACGTGTGCTC TTCCGATCTTAGAGGATCTGGGCCTCGTG ATAC | AAGTCCAG |

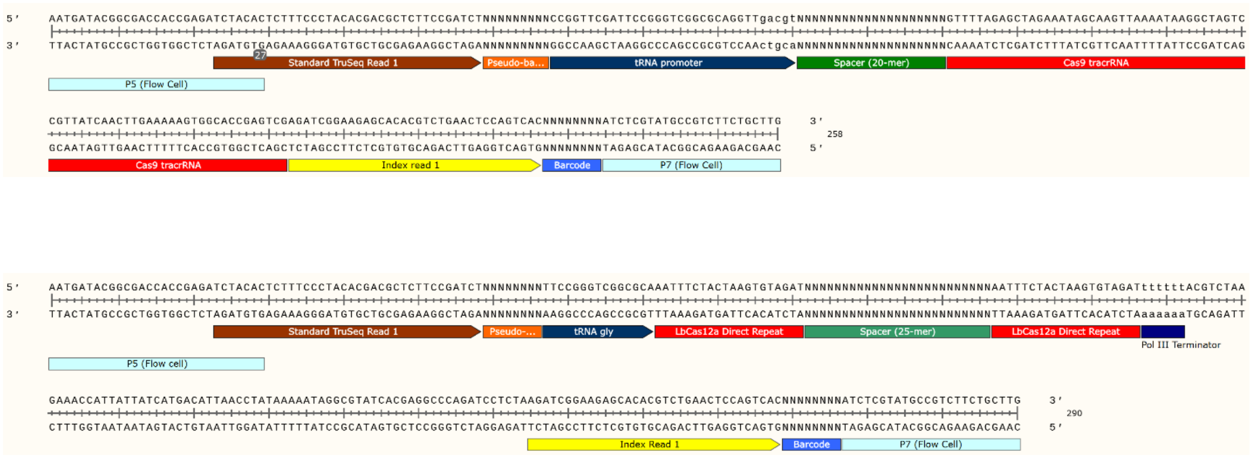


**Figure S7.** Schematic and sequence information of Cas9 (top) and Cas12a (bottom) amplicons for NGS. Amplicons contain (i) P5 and P7 sequences (light blue) that are necessary for binding with the flow cell in Illumina sequencers, (ii) TruSeq adapter (brown) for binding of the sequencing primer, (iii) a portion of tRNA$^{gly}$ (black) expressing the sgRNA, (iv) Cas9 or Cas12 spacer (green) (v) Cas12a associated direct repeats or a portion of the Cas9 tracrRNA sequence (red), (vi) Universal 8 bp Illumina barcodes (blue), (vii) Index read 1 sequence for the binding of primers to sequence the Illumina barcodes, and (viii) 4-9 nt pseudo-barcodes (orange) at the 5' end between the TruSeq and tRNA$^{gly}$ which help demultiplex replicates that contain the same illumine barcode.

**Table S11.** Parameters for bioinformatics tools used in analysis of NGS reads

| Tool | Version | Parameters* |
|---|---|---|
| FastQC | v0.11.8 | Default settings |
| Cutadapt | Galaxy Version 1.16.6 [4] | The 3 biological replicates of a given sample at a given time-point always had the same reverse primer containing the Illumina barcode, and forward primers ILU1-F, ILU3-F and ILU4-F; or ILU2-F, ILU3-F and ILU4-F each containing different pseudo-barcodes. Thus Cutadapt was used to demultiplex biological replicates from each other.<br>▪ 5' (Front) anchored 6 bp pseudo-barcodes to be demultiplexed (-g): ^NNNNNN (refer to previous table for pseudo-barcode-forward primer association).<br>▪ Maximum error rate (--error-rate): 0.2<br>▪ Match times (--times): 1<br>▪ Minimum overlap length (--overlap): 4<br>▪ Multiple output: Yes (Each demultiplexed readset is written to a separate file) |
| Trimmomatic | v0.38 | ▪ HEADCROP: 29 (if amplified by ILU1-F); or 31 (if amplified by ILU2-F); or 32 (if amplified by ILU3-F); or 34 (if amplified by ILU4-F)<br>▪ CROP: 25 |
| Bowtie2 | v2.4.2 | ▪ Number of allowed mismatches in seed alignment (-N): 1<br>▪ Length of the seed substring (-L): 21<br>▪ Function governing interval between seed substrings in multiseed alignment (-i): S,1,0.50<br>▪ Function governing maximum number of ambiguous characters (--n-ceil): L,0,0.15<br>▪ Alignment mode: end-to-end<br>▪ Number of attempts of consecutive seed extension events (-D): 20<br>▪ Number of times re-seeding occurs for repetitive reads: 3<br>▪ Save mapping statistics: Yes |

Note: All parameters other than those mentioned here are kept at default values.

**Table S12.** Correlation of SRA files names to demultiplexing information

| SRA file name | SRA sample name | Demultiplexing needed | Pseudo-Barcode for Demultiplexing with CutAdapt** | Readsets contained |
|---|---|---|---|---|
| **GW-Cpf1_Control-2_S2_R1_001.fastq.gz** | PO1f_dku70_day2_All3reps | Yes | ^AGATCG | Replicate #1 |
| | | | ^GCTATT | Replicate #2 |
| | | | ^CAGGAC | Replicate #3 |
| **GW-Cpf1_Control-4_S4_R1_001.fastq.gz** | PO1f_dku70_day4_All3reps | Yes | ^AGATCG | Replicate #1 |
| | | | ^GCTATT | Replicate #2 |
| | | | ^CAGGAC | Replicate #3 |
| **GW-Cpf1_Control-6_S6_R1_001.fastq.gz** | PO1f_dku70_day6_All3reps | Yes | ^AGATCG | Replicate #1 |
| | | | ^GCTATT | Replicate #2 |
| | | | ^CAGGAC | Replicate #3 |
| **Yl-Cpf1_CS-2_S2_R1_001.fastq.gz** | PO1f_LbCas12a_dku70_day2_All3reps | Yes | ^AGATCG | Replicate #1 |
| | | | ^GCTATT | Replicate #2 |

| | | | | |
|---|---|---|---|---|
| | | | ^CAGGAC | Replicate #3 |
| **Yl-Cpf1_CS-4_S4_R1_001.fastq.gz** | PO1f_LbCas12a_dku70_day4_All3reps | Yes | ^AGATCG | Replicate #1 |
| | | | ^GCTATT | Replicate #2 |
| | | | ^CAGGAC | Replicate #3 |
| **Yl-Cpf1_CS-6_S6_R1_001.fastq.gz** | PO1f_LbCas12a_dku70_day6_All3reps | Yes | ^AGATCG | Replicate #1 |
| | | | ^GCTATT | Replicate #2 |
| | | | ^CAGGAC | Replicate #3 |
| **GW_Yl_Cpf1-7_S7_R1_001.fastq.gz** | LbCas12a_Library_Rep1 | No | N/A | Replicate #1 |
| **GW_Yl_Cpf1-8_S8_R1_001.fastq.gz** | LbCas12a_Library_Rep2 | No | N/A | Replicate #2 |
| **GW_Yl_Cpf1-9_S9_R1_001.fastq.gz** | LbCas12a_Library_Rep3 | No | N/A | Replicate #3 |

** The symbol '^' before the barcode sequence represents that it is anchored, i.e the read begins with the barcode sequence from the 5' end. This information is needed for demultiplexing with CutAdapt.
